# Genomic dissection of Systemic Lupus Erythematosus: Distinct Susceptibility, Activity and Severity Signatures

**DOI:** 10.1101/255109

**Authors:** Nikolaos I. Panousis, George Bertsias, Halit Ongen, Irini Gergianaki, Maria Tektonidou, Maria Trachana, Luciana Romano-Palumbo, Deborah Bielser, Cedric Howald, Cristina Pamfil, Antonis Fanouriakis, Despoina Kosmara, Argyro Repa, Prodromos Sidiropoulos, Emmanouil T. Dermitzakis, Dimitrios T. Boumpas

## Abstract

Recent genetic and genomics approaches have yielded novel insights in the pathogenesis of Systemic Lupus Erythematosus (SLE) but the diagnosis, monitoring and treatment still remain largely empirical^1,2^. We reasoned that molecular characterization of SLE by whole blood transcriptomics may facilitate early diagnosis and personalized therapy. To this end, we analyzed genotypes and RNA-seq in 142 patients and 58 matched healthy individuals to define the global transcriptional signature of SLE. By controlling for the estimated proportions of circulating immune cell types, we show that the Interferon (IFN) and p53 pathways are robustly expressed. We also report cell-specific, disease-dependent regulation of gene expression and define a core/susceptibility and a flare/activity disease expression signature, with oxidative phosphorylation, ribosome regulation and cell cycle pathways being enriched in lupus flares. Using these data, we define a novel index of disease activity/severity by combining the validated Systemic Lupus Erythematosus Disease Activity Index (SLEDAI)^1^ with a new variable derived from principal component analysis (PCA) of RNA-seq data. We also delineate unique signatures across disease endo-phenotypes whereby active nephritis exhibits the most extensive changes in transcriptome, including prominent drugable signatures such as granulocyte and plasmablast/plasma cell activation. The substantial differences in gene expression between SLE and healthy individuals enables the classification of disease versus healthy status with median sensitivity and specificity of 83% and 100%, respectively. We explored the genetic regulation of blood transcriptome in SLE and found 3142 *cis*-expression quantitative trait loci (eQTLs). By integration of SLE genome-wide association study (GWAS) signals and eQTLs from 44 tissues from the Genotype-Tissue Expression (GTEx) consortium, we demonstrate that the genetic causality of SLE arises from multiple tissues with the top causal tissue being the liver, followed by brain basal ganglia, adrenal gland and whole blood. Collectively, our study defines distinct susceptibility and activity/severity signatures in SLE that may facilitate diagnosis, monitoring, and personalized therapy.

---

Systemic Lupus Erythematosus (SLE) is the prototypic systemic autoimmune disease that manifests a wide range of clinical and molecular abnormalities^2,3^. Despite advances in the pathogenesis and treatment, several unmet needs exist. To cite a few, the molecular events explaining the variability of SLE and the interspersing periods of inactivity and activity remain unspecified. The disease is often challenging to diagnose, especially at early stages, and there is lack of robust diagnostic criteria^4^. Existing instruments to monitor SLE suffer from inherent limitations and there are no accurate biomarkers of disease activity and severity. Moreover, the extent to which clinically-defined therapeutic targets correlate with changes in transcriptome is not known. Notwithstanding the successful introduction of the first targeted biological agent in SLE^5^, a considerable proportion of patients will be unresponsive to existing treatments, highlighting the need for novel, targeted therapies based on the underlying immune aberrancies. To examine these, we performed transcriptome profiling by RNA-seq (**Supplementary Figure 1**) in 142 SLE patients (**Supplementary Table 1**) with varying levels of disease activity, and compared it to matched healthy individuals.

Our data show widespread transcriptome perturbations in SLE with 6730 differentially expressed genes (DEGs) (false discovery rate [FDR] 5%) (**Supplementary Figure 2A** and **Supplementary Table 2**); 3977 genes were upregulated (59.1%) and 2753 genes were downregulated (40.9%) in SLE versus healthy individuals. The enriched KEGG pathways^6^ (5% FDR) for DEGs are shown in **Supplementary Figures 2B-D** with novel and previously identified pathways implicated in SLE being confirmed, such as the IFN signature represented in the *Herpes simplex virus (HSV)* and the *NOD-like receptor signaling* pathways^7^. A broad view of the biological regulation of SLE is provided by a network enrichment map of significantly enriched GO terms derived from DEGs (**Supplementary Figure 2E**). These results reveal marked aberrancies and specific signatures in SLE whole blood transcriptome.

Whole blood assays, while relevant to define complex inflammatory signatures^8,9^, do not inform on cell-specific mechanisms. To address this, we estimated the proportions of different immune cell types for each individual by using blood transcriptome deconvolution implemented in CIBERSORT^10^. We report significant differences in blood cell composition including reduction of naïve B-cells^11^ and natural killer cells^12,13^, and increase of memory activated CD4^+^ T-cells^14^ and myeloid-linage cells^15-18^ in SLE versus healthy individuals (**Figure 1A**). Next, we defined a global gene expression signature independent of the differences in cell composition by controlling for the estimated proportions of cell types, and found 1613 DEGs (5% FDR) between SLE and healthy individuals, which represents a 76% decrease in DEGs from the previous analysis. Pathway and GO term enrichment analysis revealed that the IFN signature (**Figure 1B-C**) is independent of cell type composition. By interrogating the interferome database^19^, we found that the DEGs are indicative of both type I and type II IFN response, while they also showed enrichment in the *p53 signaling* pathway (**Figure 1B**), which is implicated in apoptotic cell clearance and maintenance of immune tolerance^20,21^. These data demonstrate a robust transcriptional signal – independent of blood cell type composition – that could facilitate the discovery of novel biomarkers.

**Figure 1:**
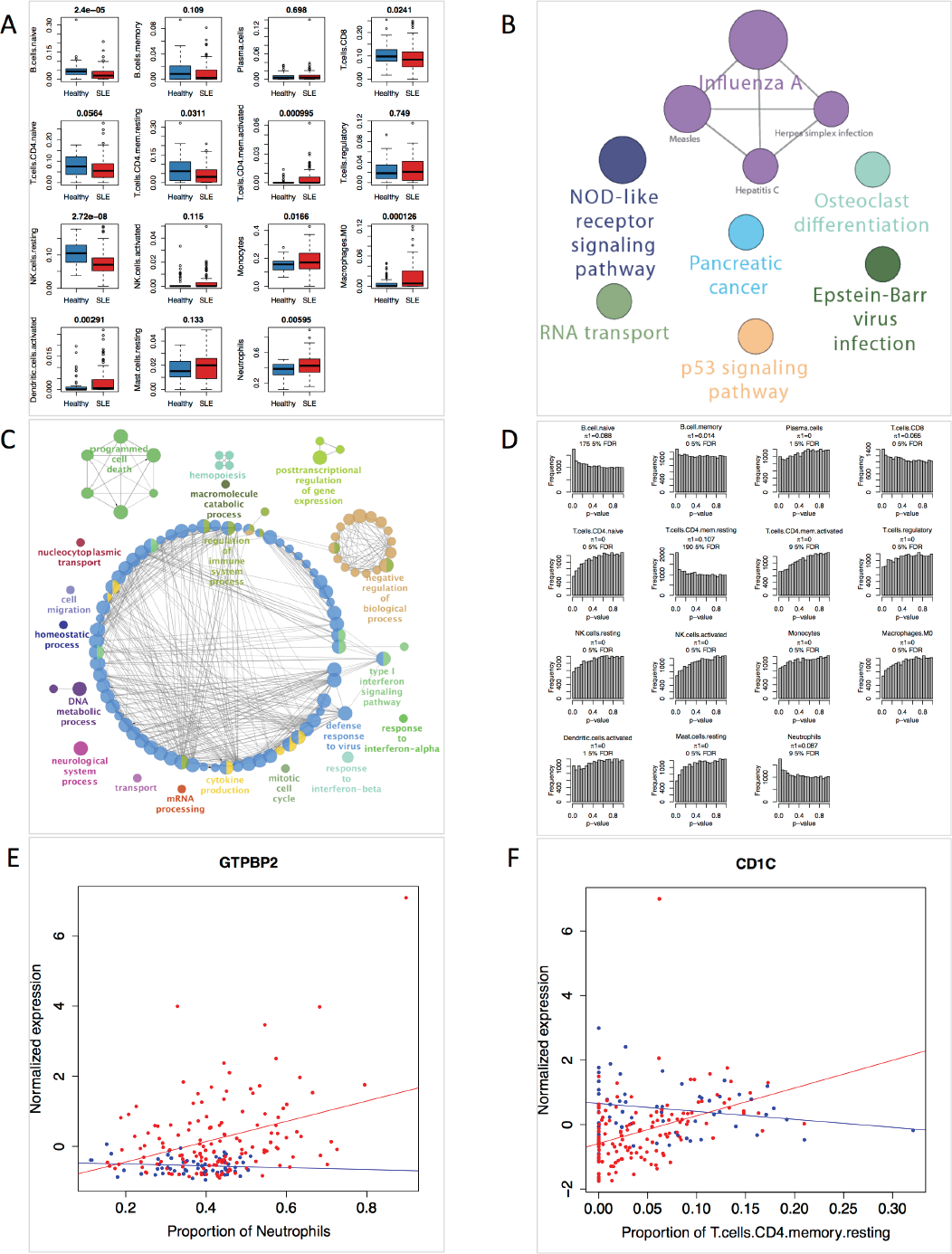
**(A)** Estimated proportions of different immune cell counts in healthy and SLE individuals. For every cell type the Mann-Whitney Wilcoxon test p-value comparing healthy and SLE patients is displayed on top. **(B)** Pathway enrichment analysis with all the DEGs after correcting for cell type proportion estimates. **(C)** Mechanistic map of the biological regulation in SLE after correcting for cell type proportion estimates. There is a prevalent type I IFN regulation independent of immune cell type composition. **(D)** Histogram’s of p-values for the interaction term (disease X estimated proportions) revealing cell type specific effects for SLE. For every cell type the proportion of estimated true positives (π1) and the number of significant genes at 5% FDR is presented. **(E)** Disease by estimated Neutrophils proportion interaction for the gene *GTPBP2*. X-axis indicate the estimated proportion of Neutrophils while y-axis indicate the normalized expression. Red dots indicate SLE patients while blue dots indicate healthy individuals. **(F)** Disease by estimated T-cell CD4 memory resting proportion interaction for the gene *CD1C*. X-axis indicate the estimated proportion of Neutrophils while y-axis indicate the normalized expression. Red dots indicate SLE patients while blue dots indicate healthy individuals.

To unravel cell-specific gene perturbations, we examined whether cell type composition has a different effect on gene expression between the SLE and healthy state. We controlled for the estimated proportions of all other immune cell types and tested for the significance of the disease × estimated cell proportion interaction term. We quantified the proportion of true positives estimated from the enrichment of significant p-values (π1)^22^ (**Figure 1D**). For naive and memory B-cells, CD4+ memory resting T-cells, CD8+ T cells, and neutrophils, the corresponding π1 values were above zero, suggesting that varying proportion of these cells interacts with the effects that cell type proportions have on gene expression in SLE versus healthy individuals. Illustratively, increasing proportion of neutrophils correlated positively with *GTPBP2* (GTP Binding Protein 2) in SLE but not in healthy individuals (**Figure 1E**). The same trend was observed for the correlation between CD4+ memory resting T-cells and *CD1c* (cluster differentiation 1c) (**Figure 1F**). GTPBP2 interacts with IRF5 to regulate type I IFN production^23^, while CD1c is a cell membrane glycoprotein^24^ that mediates the presentation of modified peptides to CD1c-restricted T cells^25^ and has been linked to autoimmunity^26,27^. Together, differences in whole blood transcriptome in SLE are driven by both altered abundances of circulating immune cell types and cell-specific regulation of gene expression depending on disease status.

Remission of disease activity has been introduced as therapeutic target in SLE^28^ but whether this is mirrored by transcriptome changes remains unknown. This has implications for predicting the risk for subsequent flare and assessing the need for continuing long-term immunosuppression. To this end, we determined a ‘core’ gene expression signature that persists in the absence of disease activity following treatment. First, we applied PCA on the transcriptome in clinically active and inactive patients and healthy individuals (**Supplementary Table 1** and **Figures 2A-C**). Inactive SLE were clearly differentiated from healthy **(Figure 2B)** but not from active SLE individuals **(Figure 2C)**, signifying persistently deregulated gene expression despite disease remission. We took the intersection of DEGs in healthy versus active SLE (4938 DEGs 5% FDR; **Supplementary Table 3**) and healthy versus inactive SLE (4658 DEGs; **Supplementary Table 4**) that are not DEGs in active versus inactive SLE (377 DEGs; **Supplementary Table 5**), to reach 2726 DEGs which comprise the ‘core’ disease gene signature. These genes were enriched in biological processes and GO terms related to the *regulation and response of the immune system* (**Supplementary Figure 3**) suggesting persistence of immune system activation and inflammation despite remission of clinical signs and symptoms. By comparing patients with clinically inactive SLE but evidence for serologic activity (high anti-dsDNA titers, low serum complement)^29^ against those who are both clinically and serologically inactive, we found no DEGs at 5% FDR, corroborating studies showing similar favorable prognosis for these two groups^30^.

**Figure 2:**
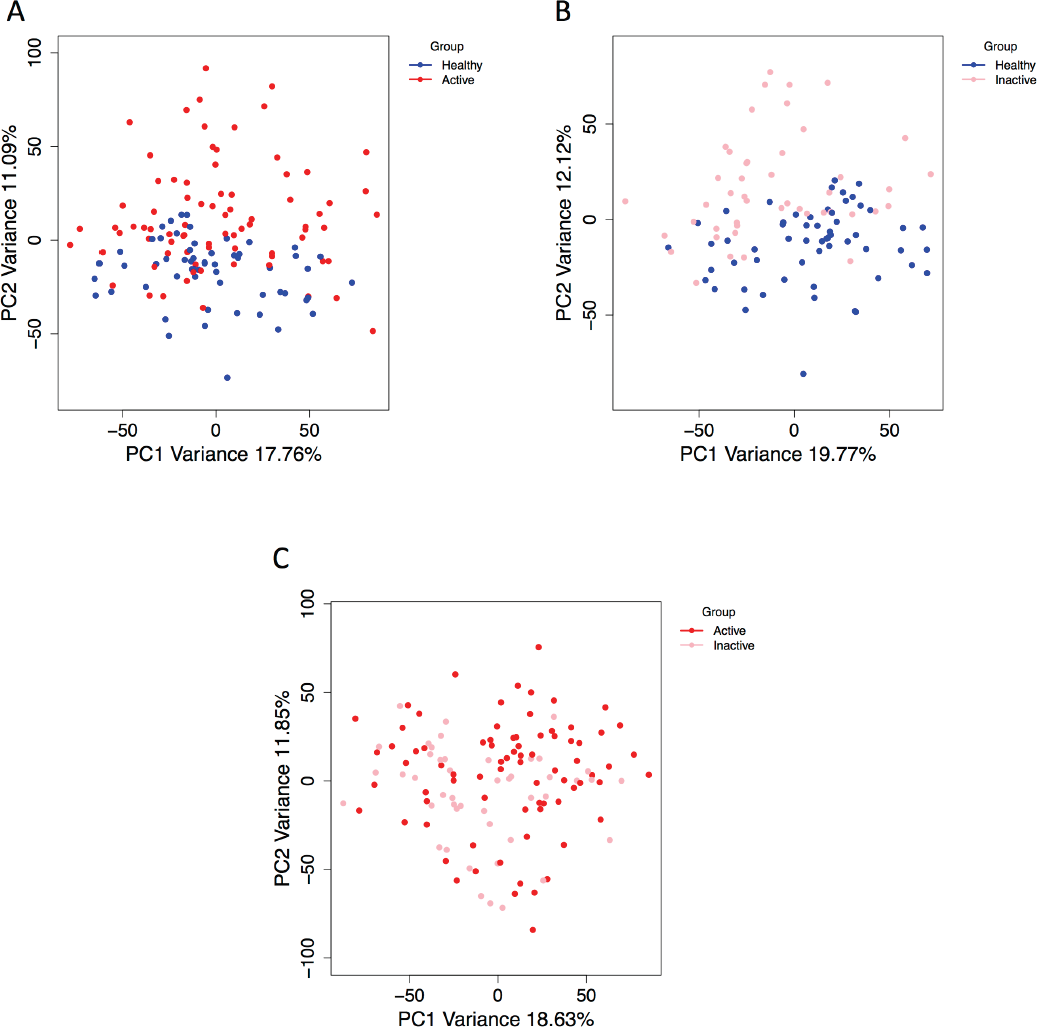
**(A)** Principal Component Analysis (PCA) of whole transcriptome between healthy and Active SLE individuals. The two first Principal Components are plotted. PC2 is clearly differentiating the 2 groups implying differences in gene expression. **(B)** PCA analysis between healthy and Inactive SLE individuals. PC2 is clearly differentiating the 2 groups indicating that even in remission the transcriptome of SLE patients is different compared to healthy individuals. **(C)** PCA analysis between Active and Inactive SLE patients. The first 2 PCs do not differentiate the 2 groups suggesting that there are not large differences in gene expression between them.

Flares of disease activity are frequent in SLE and contribute to accrual of irreversible organ damage^31^. Defining the signature of increased SLE activity has pathogenic and clinical ramifications. To this end, we selected the DEGs from the comparison of inactive versus active SLE that are not included in the ‘core’ disease signature. A total 365 DEGs were identified, which were enriched for KEGG pathways such as oxidative phosphorylation, consistent with the described alterations in mitochondrial mass and membrane potential in lupus T cells^32-34^ and the enhanced oxidative stress^35-37^. Other enriched pathways included ribosomes and cell cycle. Thus, increases in SLE activity may be linked to perturbed expression of genes that regulate metabolism, protein synthesis and proliferation of peripheral blood immune cells. The flare gene signature could enable the earlier identification of an impending clinical flare and more effective use of pre-emptive treatment, an unmet need in SLE in view of the limitations of traditional serologic tests^38,39^.

Biomarkers that accurately reflect disease severity and the underlying molecular pathogenesis, represent an additional unmet need in SLE. We assessed transcriptome differences according to varying degrees of disease activity/severity by using the validated SLEDAI-2K index as a quantitative measure (0=inactive; 1–5=mild; 6–10=moderate; >10=severe disease)^40^, among patients and healthy individuals (fixed score of -1). We performed PCA (**Figure 3A**) on the identified 3690 DEGs (5% FDR) (**Supplementary Table 6**) and defined PC1 (explaining 23% of the variance) as a new variable summarizing the expression properties of genes that recapitulate SLE severity (**Figure 3B**). Notably, PC1 clusters closely patients with inactive and low disease activity, which is consistent with evidence that both these states have a favorable outcome^41,42^. Pathway analysis showed enrichment in the *oxidative phosphorylation* and *cell cycle* KEGG pathways, similar to the ‘flare’ signature, suggesting that these two biological processes may be implicated not only in disease activity but also in severity. Functionally grouped networks of GO terms revealed gene signatures related to protein ubiquitination, electron transport chain, protein phosphorylation, cell cycle, defense response and regulation of response to stress (**Figure 3C**), all of which have been linked to SLE^37,43,44^. These results point out the involvement of multiple molecular pathways and biological processes in determining SLE progression/severity and suggest that whole blood transcriptome may serve as a robust biomarker of SLE activity and severity.

**Figure 3:**
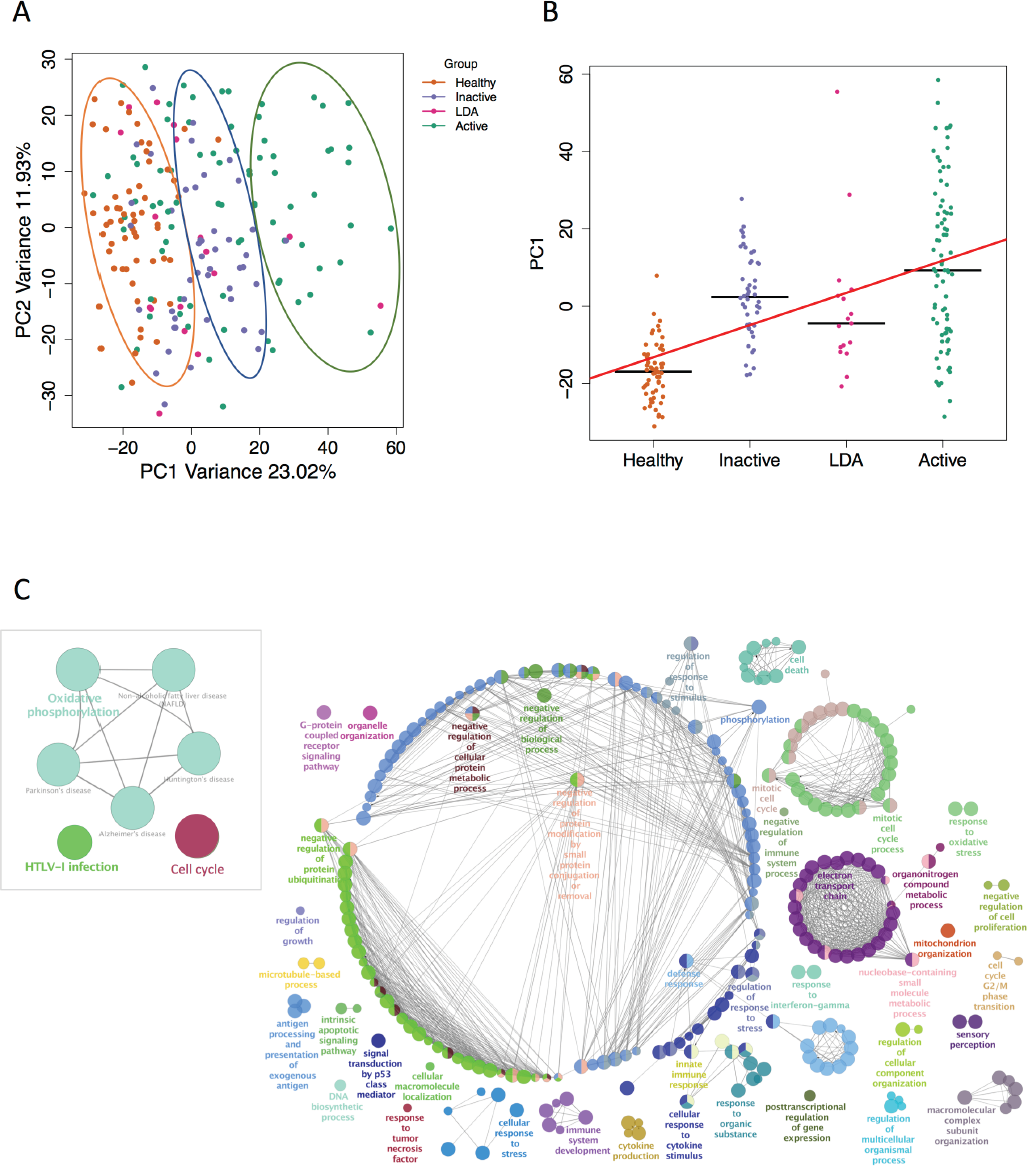
**(A)** Principal Component Analysis (PCA) of 3690 DEGs found by using SLEDAI score as a quantitative measurement of SLE severity. The first two Principal Components are plotted. PC1 is capturing the progression of activity/severity of SLE since is separating the different groups of activity. **(B)** Jitter plot of PC1 weights. For each group, the median is plotted. The regression line is plotted with red (p-value5.86e-17). PC1 defines a new phenotype that captures SLE activity. **(C)** Pathway enrichment analysis of 3690 DEGs on the left side of the plot. Oxidative phosphorylation and cell cycle KEGG pathways, that were regulated by genes belonging to the “flare” gene signature is also enriched suggesting that these genes not only capture the inactive to active progression but also the different levels of severity. Functionally grouped networks of GO terms on the right. Multiple biological aberrations are captured by differences in gene expression based on different levels of SLE activity.

SLE can affect multiple organs but the molecular basis of this heterogeneity remains elusive. Accordingly, SLE patients were grouped according to predominant organ activity (**Table 1**) and multiple comparisons were carried out. By comparing patients with active renal disease (nephritis) (group 1) versus those with activity from other organs (combined groups 2,3,4), we found 136 DEGs (5% FDR). These genes were enriched in functionally grouped networks of *granulocyte activation* and *antimicrobial humoral response* (**Supplementary Figure 4**), consistent with the role of neutrophil activation^45,46^ and their death by formation of chromatin extracellular traps,^47-49^ and of plasmablasts/plasma-cells^11,50-52^ in lupus nephritis. To further discern the transcriptome basis for kidney involvement in SLE, we took the intersection of DEGs in active nephritis (group 1) versus inactive SLE (combined groups 5 and 6) (1375 DEGs, 5% FDR) with those in active versus inactive SLE (377 DEGs). A total 305 genes were common in the two comparisons, suggesting a step-wise progression of transcriptome alterations from inactive to active non-renal and active renal SLE status. By comparing patients with active versus inactive nephritis, we found global gene expression differences captured by PCA (**Supplementary Figure 5**), while differential gene expression analysis between these two groups revealed 1496 DEGs. Together, lupus nephritis displays marked and gradually enriched changes in whole blood gene expression compared to other SLE subsets, some of which may be drugable.

**Table 1:**
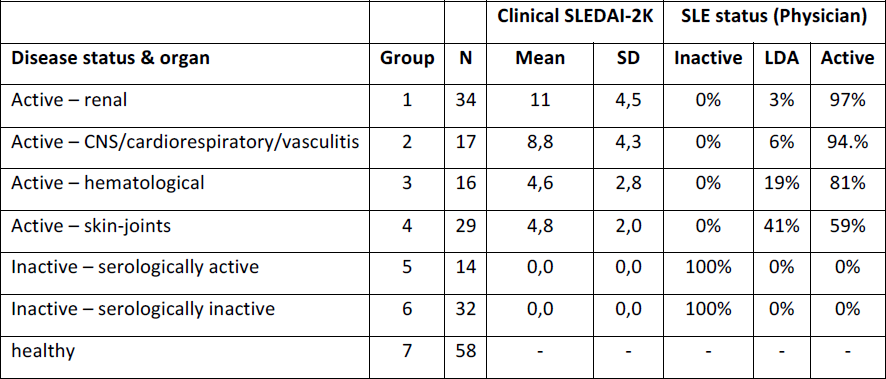
Groups of individuals based on the tissue/organ that the disease activity originates.

SLE exhibits a striking gender bias with females being affected 7 to 12 times more frequently than men, yet the latter suffering from more severe disease^53^. To gain insights into the molecular basis of this sexual dimorphism, we took the non-overlapping sets of DEGs in male versus female SLE (Bonferroni significant) and male versus female healthy (90% FDR) individuals. Six genes showed gender-biased expression in SLE (**Supplementary Figure 6A**), two of which (*SMC1A*, *ARSD*) are located on X chromosome and escape X-inactivation. *SMC1A*, encoding for the cohesin complex protein Structural Maintenance Of Chromosomes 1A^54-56^, demonstrated the strongest gender bias (**Supplementary Figure 6A-B**), and this was confirmed in purified CD4+ T-cells (**Supplementary Figure 6C**). Although preliminary, these results provide candidate genes for further studies on the sexual dimorphism in the disease.

SLE diagnosis can be challenging especially at early stages before an adequate number of features accumulate^4^. We asked whether we could accurately classify individuals based on their transcriptional profile by building multiple classifiers based on Linear Discriminant Analysis (LDA) by the use of DEGs as features (*see Methods*). We measured a median diagnostic accuracy of 90% in the validation set with 86% sensitivity and 100% specificity (**Supplementary Figure 7**). Patients who were most frequently misclassified as healthy individuals had lower frequency of renal involvement, ANA/anti-dsDNA autoantibodies, and were less frequently treated with immunosuppressive agents, suggesting an overrepresentation of milder disease. Thus, whole blood transcriptome might be used to assist the diagnosis of SLE, a finding that needs to be confirmed and validated in longitudinal studies of patients with both early and established disease.

Definition of regulatory genetic effects is crucial to our understanding of the molecular basis of complex diseases^57^. We explored the genetic regulation of blood transcriptome in SLE by assessing expression-Quantitative Trait Loci (eQTLs) in our cohort. We found 3142 *cis*-eQTLs (5% FDR), with highly significant cis-eQTLs clustering close to the transcription start site of the genes (**Supplementary Figure 8A-B**). These eQTLs replicated well (pi=0.89, **Supplementary Figure 8C**) with a blood RNA-seq study in healthy donors^58^, suggesting the lack of disease specificity. This could be due to lack of adequate power to detect such effects and/or the need to study specific immune cell types. Assessing eQTLs from multiple tissues could enhance our interpretation of GWAS signals, as most of them reside in non-coding genome areas^59^ and enable the identification of the causal genes and tissues. To estimate the tissue(s) that determine the genetic causality of SLE, we employed the Regulatory Trait Concordance (RTC) method^57,60^ that estimates the causal tissues by using eQTLs in 44 tissues from the GTEx consortium^61^ (**Figure 4**). We found that the top causal tissue is liver followed by brain basal ganglia, adrenal gland and whole blood, suggesting that the genetic causality of SLE arises from multiple tissues. The finding of liver being linked as the top causal tissue is in agreement with our result that SLE exacerbation is associated with changes in expression of genes that regulate metabolism^62^.

**Figure 4:**
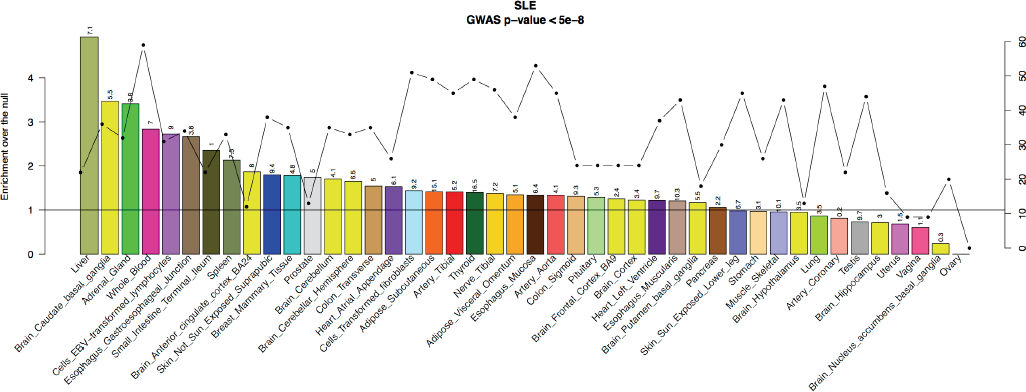
**(A)** Estimation of the genetic causality of SLE in 44 tissues from GTEx. On the primary y-axis, the enrichments over the null per tissue are plotted as bars and on the secondary y-axis number of GWAS variants that co-localized with eQTLs per tissue are plotted as a line. The horizontal black line indicates the null. On top of each of the bars are the -log10 Benjamini-Hochberg corrected p-values for the enrichments.

In summary, SLE is an autoimmune disease with marked clinical and immunological heterogeneity, waxing and waning course, and variable response to immunosuppressive and biological treatments. In a previous study, Banchereau *et al* ^63^ profiled the blood transcriptome by microarrays in a longitudinal cohort of pediatric patients. However, their analysis focused on detection of immune correlates of disease activity and treatment effects based on *a priori* defined modules of transcripts co-expressed in blood across various immunological conditions. Our RNA-seq analysis in a well-characterized cohort of adult SLE patients provides an unbiased characterization of whole-blood gene signatures associated not only with susceptibility but also, with clinically relevant disease progression such as increased activity and major organ involvement. We also defined cell-specific effects of the disease on gene expression, which may provide insights into the pathogenic role of specific immune cell types by the further studies in purified cells. DEGs are involved in biological networks and pathways that are pertinent to SLE pathogenesis and may represent putative therapeutic targets. Importantly, we show that whole blood transcriptome can classify SLE versus healthy individuals with excellent specificity putting forward the possibility of using blood gene profiling as a diagnostic tool. Finally, our studies on the genetic variation on gene expression suggest that in addition to whole blood, other tissues such as the liver may determine genetic causality in SLE.

## Acknowledgments

This work was funded by the Foundation for Research in Rheumatology (FOREUM) and in part by the Greek General Secretariat of Research and Technology ‘Aristeia’ action of the Operational Program ‘Education and Lifelong Learning’ (co-funded by the European Social Fund and National Resources, Aristeia I 2344 to DB) and the European Research Council (ERC) Advanced Grant (“Lupuscare” to DB). GB was supported by the Greek State Scholarships Foundation-ΙKY (Postdoc Fellowships of Excellence-Siemens) and the European Union Seventh Framework Programme (project TransPOT; FP7-REGPOT-2011-1).

## Author contributions

NIP carried out the analyses. NIP and GB drafted the manuscript, with contributions from all authors. GB recruited and took care of patients, collected blood samples, extracted RNA and DNA. HO performed the RTC analysis. LRP and DB prepared the RNA-seq libraries and CH processed the RNA-seq and genotyping data. IG contributed in recruitment of patients and healthy individuals, and extracted clinical data from the medical charts. DK isolated CD4+ T-cells from peripheral blood samples and performed RT-PCR studies. MT, MT, CP, AF, AR and PS contributed patient samples and participated in the analyses of data. DTB and ETD conceived the study. All authors read and approved the final manuscript for publication.

## Methods

### Patients, samples collection and clinical assessment

SLE patients, diagnosed according to board-certified rheumatologist’s judgment and/or the 1997 revised ACR classification criteria^64^, were enrolled from the rheumatology clinics at the University Hospital of Heraklion, University Hospital C.F.R Cluj Napoca, General Hospital of Athens ‘Laikon’, ‘Attikon’ University Hospital in Athens and ‘Hippokration’ Hospital of Thessaloniki. Demographic and clinical characteristics including history of biopsy-proven nephritis, ACR classification criteria, serum autoantibodies, assessment of disease activity (physician’s global assessment [PGA], SLEDAI-2K^40^), definitions of Lupus Low Disease Activity State and remission^41,42^, and use of medications, were evaluated by a standardized protocol. Informed consent was obtained from adults and the parents those younger than 18 years of age and all procedures were followed in accordance with protocols approved by the local institutional review boards. Patients were asked to withdraw all lupus medications for 12 hours prior to blood collection. Venipuncture was performed to collect blood in PaxGene RNA tubes (Qiagen) for mRNA extraction and in EDTA tubes for DNA extraction.

### Study design

We performed RNA-seq to measure gene expression in whole blood from SLE patients and matched for age and sex healthy individuals. After quality control (**Supplementary Figure 1**), we obtained gene quantifications for 142 SLE patients and 58 healthy volunteers, measuring the expression of ~21.000 genes. All individuals were genotyped and imputed towards the 1000 genomes phase III reference panel^65,66^, yielding a set of ~6.9 millions of variants. For all SLE individuals we collected a variety of clinical phenotypes such as the 1997 American College of Rheumatology revised criteria for classification of SLE, presence of serum autoantibodies, physician-rated disease activity (PGA), an index measuring the activity of the disease (SLEDAI-2K)^1,67^, and the medical treatment at the time of sample collection.

### Genotyping

All the individuals were genotyped with the Illumina HumanCoreExome-24 array. The individuals were phased with SHAPEIT^68^ and imputed to the 1000 Genomes Project Phase III^66^ using IMPUTE2^69^. After imputation, we filtered out SNPs with MAF <=0.05, IMPUTE score < 0.4 and Hardy-Weinberg equilibrium p-value <5e-7. After filtering we ended up with ~7M variants for association testing.

### RNA sequencing, mapping and quantifications

RNA libraries were prepared with the Illumina TruSeq sample preparation kit and were sequenced on Illumina HiSeq2000 according the manufacturer’s instructions. 49 bp paired-end reads were mapped to the GRCh37 reference human genome using the GEM mapper^70^. We kept reads with mapping quality >150 to quantify exons and genes corresponding to reads that are uniquely mapped to the genome, with the correct orientation between pairs, allowing for 5 mismatches for both reads of a pair. Exon quantification was performed using the GENCODE annotation v15^71^. The overlapping exons of a gene were merged into a meta-exon unit with starting coordinates the start position of the first exon that is merged and end position the end position of the last exon that is merged. We counted reads as mapped to this meta-exon unit if there was an overlap between either the start or end position of the read with the meta-exon. Gene level quantifications were then obtained by summing the meta-exons counts for each gene.

### Quality control of RNA sequencing data

Several metrics were used to assess the quality of the RNA-seq data (Supplementary Figure 1A). We measured the proportion of exonic/total reads in quantified exons per sample. Samples with a ratio less than 0.2 were excluded from further analysis as technical outliers. We further excluded samples being outliers in the PCA plot (Supplementary Figure 1B). These samples were also outliers based on the proportion of total reads mapped to hemoglobin genes (Supplementary Figure 1C). Two samples were also excluded from further analysis based on discrepancy of the sex identified by the RNA-seq data and our clinical records (Supplementary Figure 1D). Finally, MBV^72^ was used to correct for any sample label swaps or cross contamination for the RNA-seq and the genotyped data without identifying any swaps in our dataset.

### Differential gene expression analysis

We included 21851 genes in the analysis and used this set of genes in all downstream analyses for consistency. A gene was included if it had at least 5 reads in 10% of either the SLE or the healthy individuals. DESeq2^73^ was used to call differentially expressed genes by including GC content, RNA integrity (RIN), center of collection, insert size mode, age, gender, amount of RNA to construct the library and plate number as technical covariates. These covariates were found to influence gene expression by performing step wise regression between gene expression and all the technical covariates that were obtained from the experimental procedure. Benjamini-Hochberg was used to assess significance at 5% or 1% false discovery rate (FDR). To identify genes that were differentially expressed despite the immune cell composition we used the same covariates as before by including the estimated cell proportions as covariates in the model. To assess cell specific differentially expression we used the latter model by including a *disease* × *estimated proportion status* interaction term and obtained p-values for every gene for the interaction term. For this analysis, we estimated the proportion of true positives from the enrichment of significant p-values by using the π1 statistics. To specify the DEGs based on disease severity/activity the SLEDAI-2K index was used as a quantitative measurement by fixing the score to -1 for healthy individuals. To call up and down regulated DEGs we used Spearman rank correlation between gene expression and the SLEDAI index.

### Pathway and GO term enrichment analysis

To functionally characterize the identified DEGs a comprehensive functional enrichment gene set analysis for KEGG pathways and GO terms was performed by using the ClueGO^74^ plugin of Cytoscape^75^. Briefly, ClueGO visualize the functionally grouped networks of enriched pathways and GO terms. The enrichment is based on a two-sided hypergeometric test and corrected for multiple testing with either Benjamini-Hochberg for KEGG pathways or Bonferroni for GO terms by setting the threshold of the adjusted p-values in each case at 0.05.

We used a more stringent approach with GO terms by Bonferroni correction and fusion criteria implemented in the ClueGO plugin to reduce the plethora of the terms that have the same enriched genes allowing only for the most representative “parent” or “child” terms in the networks. Each node represents a pathway or a GO term and the size of each node represents the enrichment significance. The connection between nodes is based on Cohen’s kappa statistic score (≥ 0.4) which depends on the gene sharing between nodes. For each network, only the most significant node is labeled.

### Cell type estimation

We used CIBERSORT^10^ to estimate the proportion of different immune cell types in whole blood in our dataset. CIBERSORT uses a well-defined gene expression signature (LM22) that consists of 547 genes that precisely differentiate mature human hematopoietic cells. To perform the analysis, we used the LM22 gene matrix as reference and normalized values for the library size per sample in our dataset obtained from the *estimateSizeFactors* function in DESeq2.

### CD4+ T-cell purification and RT-PCR

Peripheral blood mononuclear cells (PBMCs) from SLE patients and age-matched healthy blood volunteers were isolated by Ficoll-Histopaque (Sigma-Aldrich) density-gradient centrifugation of heparinized venous blood. CD4+ T lymphocytes (98% purity) were isolated by negative immunomagnetic selection (Miltenyi Biotec, Bergisch Gladbach, Germany). Total RNA was extracted using the TRIzol^™^ extraction method and the Turbo DNAse kit (Ambion) was used to eliminate genomic DNA contamination. cDNA was prepared using Perfect Real time cDNA Synthesis Kit (Takara) according to manufacturer’s protocol. 200ng of RNA were used as a template for every reaction. RNAse H (2U/reaction) was added to clean the resulting cDNA from any RNA and incomplete cDNA products. PCR amplification of the resulting cDNA samples was performed using appropriate volumes of KAPA SYBR^®^ FAST Universal 2x qPCR Master Mix and specific for each gene primers at a CFX Connect^™^, Real-Time System. Total volume of each PCR reaction was 20μl. Expression was normalized to GAPDH and calculated by the change-in-threshold method [2^(-ΔΔCT)]. The following primer sets were used: (5’ → 3’): *SMC1A* forward human: CAT CAA AGC TCG TAA CTT CCT CG; *SMC1A* reverse human: CCC CAG AAC GAC TAA TCT CTT CA; *GAPDH* forward human: CAT GTT CCA ATA TGA TTC CAC C; *GAPDH* reverse human: GAT GGG ATT TCC ATT GAT GAC.

### Classification

we performed Linear Discriminant Analysis (LDA) by using DEGs as features. We divided our dataset to training (90%, corresponding to 128 SLE patients and 52 healthy individuals) and validation set (10%, corresponding to 14 SLE patients and 6 healthy individuals) and run 1000 iterations. For each iteration, we performed differential gene expression analysis between SLE and healthy using the training set and built the LDA classifier based on the identified DEGs in order to measure the accuracy, sensitivity (the probability of calling an SLE individual as patient) and specificity (the probability of calling a healthy individual as healthy) of the classifier. We found a median of 6126 DEGs ranging from 4671 to 7222 DEGs. We chose the strategy of building multiple classifiers to account for the SLE heterogeneity by inserting perturbations in the models by sampling in each iteration different individuals. In doing so, we achieved high levels of accurate classification.

### eQTL mapping

eQTL analysis was performed in a 1MB window upstream or downstream the transcription start site of the gene using QTLtools^76^. We scaled the total number of raw reads per sample to the median number of all samples. We removed genes that were not expressed in 90% of the individuals and then we applied PCA to account for hidden technical variation. Significance was assessed by using the qvalue R package^22^ on beta approximated empirical p-values from 1000 permutations. Replication was assessed by using the π1 statistic on the p-value distribution obtained from significant cis-eQTLs.

### Regulatory Trait Concordance (RTC)

We have previously described RTC score to assess whether a GWAS variant and an eQTL are tagging the same causal variant^57^. Briefly, we expect that if a GWAS variant and an eQTL do tag the same causal variant, by removing the genetic effect of the GWAS variant will have a significant consequence on the eQTL association. Following up, we can now estimate the causal tissues for complex traits and diseases by measuring the tissue sharing probabilities of eQTLs and calculating probabilities that a GWAS variant and the eQTL do tag the same functional effect. By normalizing the GWAS-eQTL probabilities with the tissue sharing estimates of the eQTLs, we can estimate the tissues from which GWAS genetic causality arises^60^.

## Supplementary information

**Supplementary Table 1.**
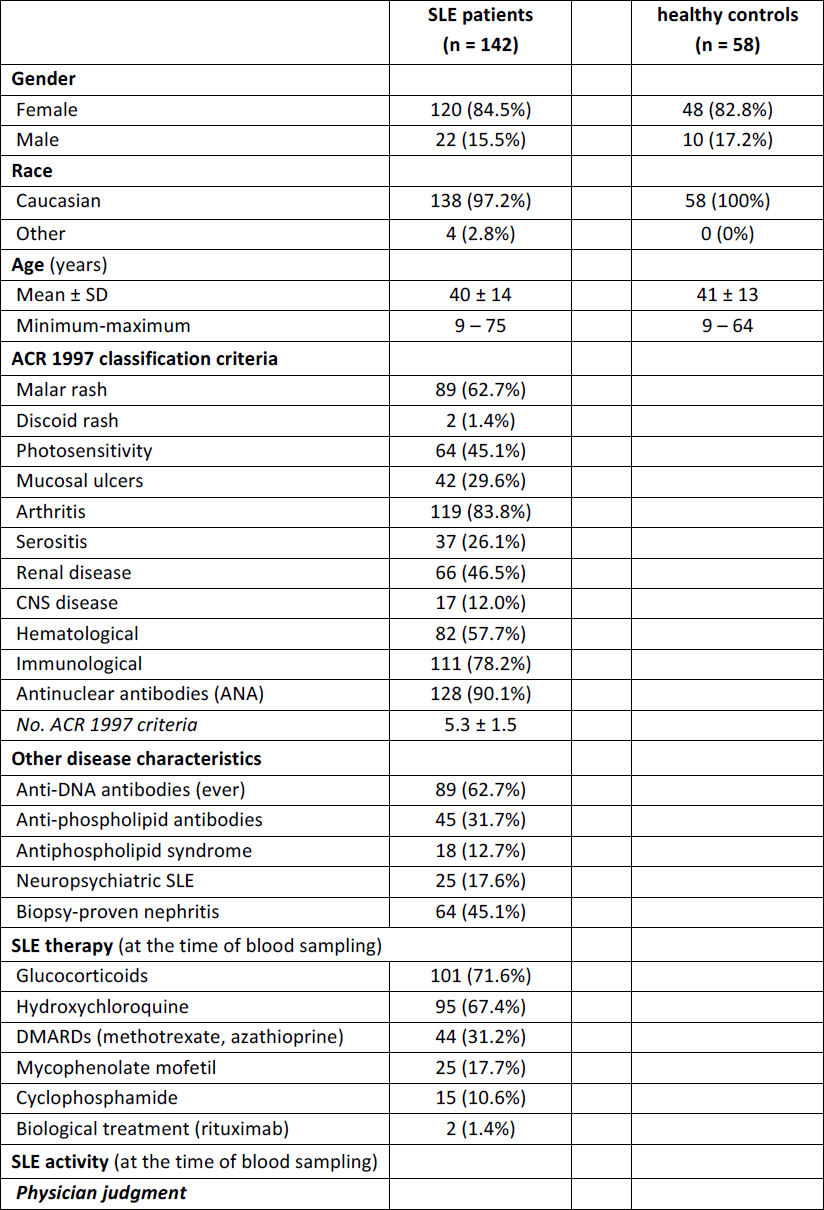
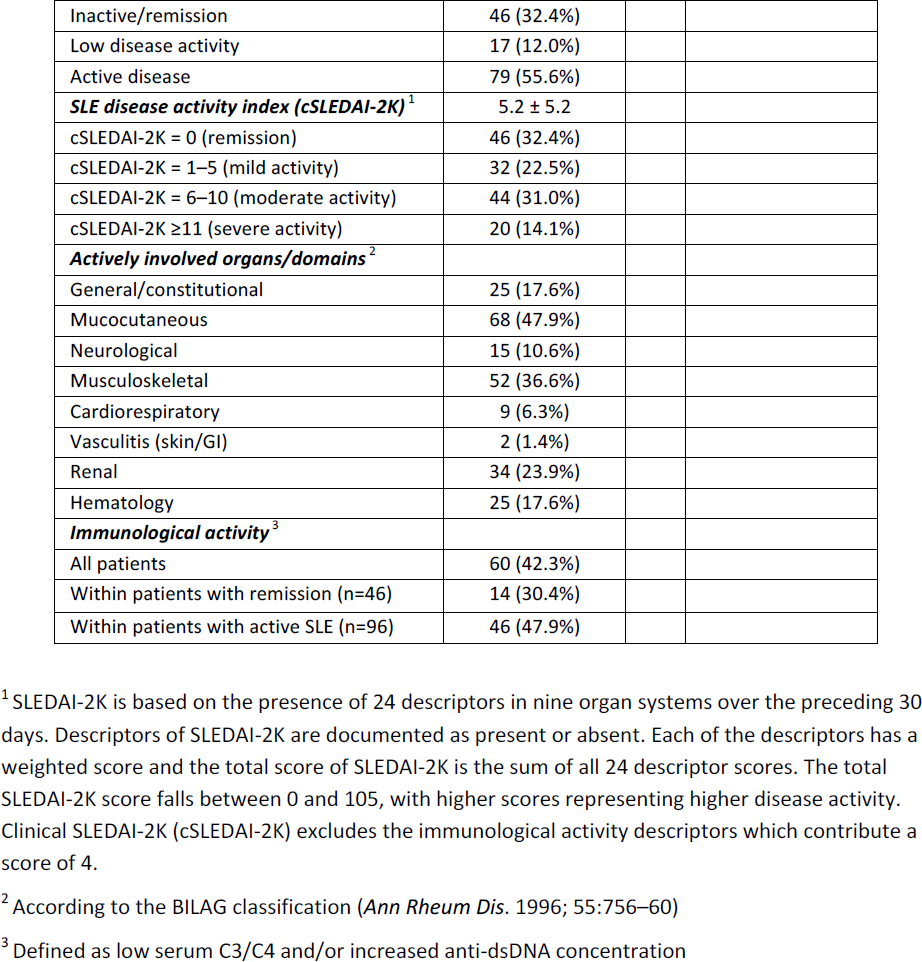
Demographic and clinical characteristics of SLE patients and healthy individuals.

**Supplementary Figure 1:**
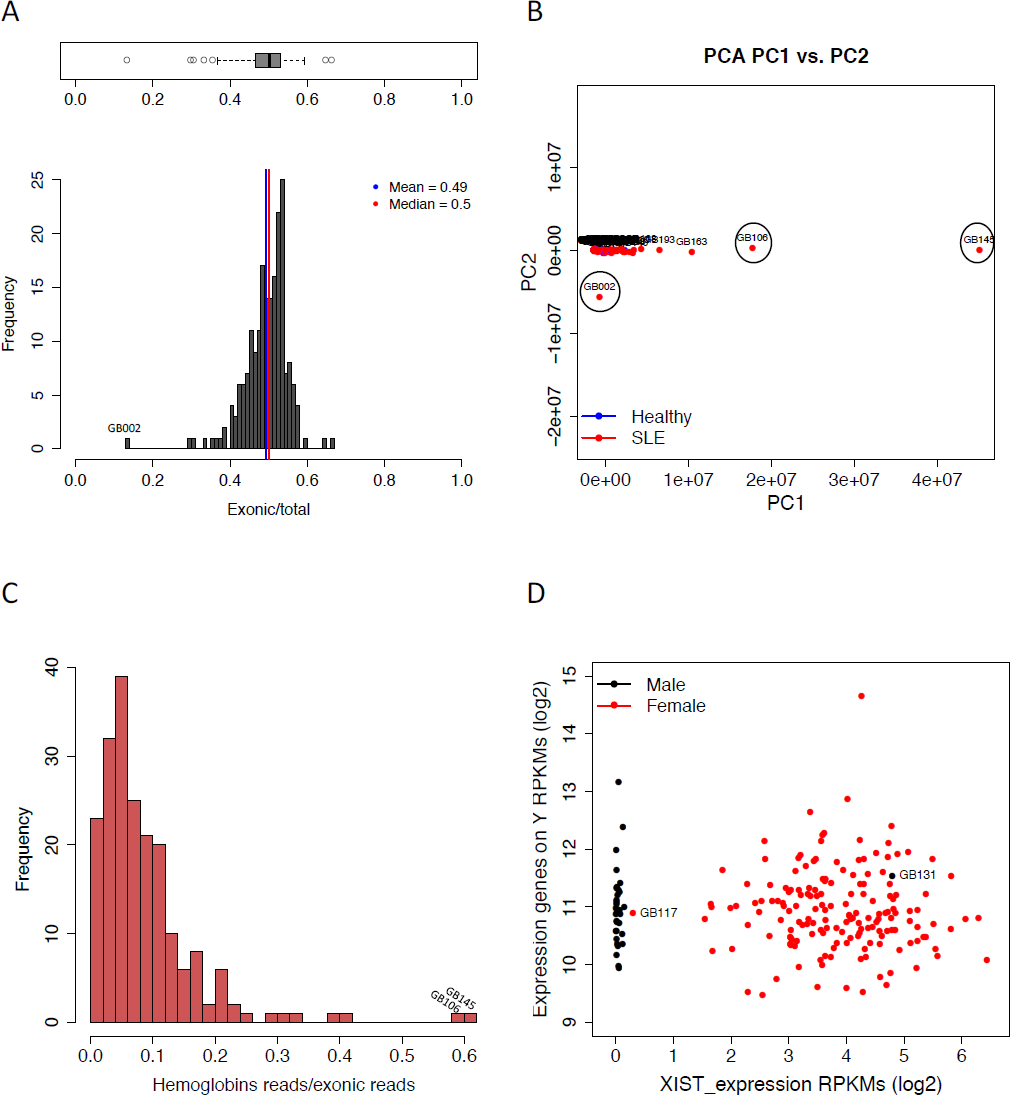
(**A**) Fraction of reads mapping to exons per sample. The vertical red line indicates the median and the blue the mean. Samples with less than 0.2 exonic/total reads were considered as technical outliers. (**B**) PCA analysis of gene expression of all the samples. Samples in circles were considered as technical outliers. **(C)** Proportion of reads mapped to hemoglobin genes. Samples above 0.5 were considered as technical outliers (same samples as B). (**D)** Sex specific expression, in x-axis RPKMs (log2) of the female specific *XIST* gene versus the sum of RPKMS (log2) of the genes in Y chromosome excluding genes mapped in the pseudo-autosomal region.

**Supplementary Figure 2:**
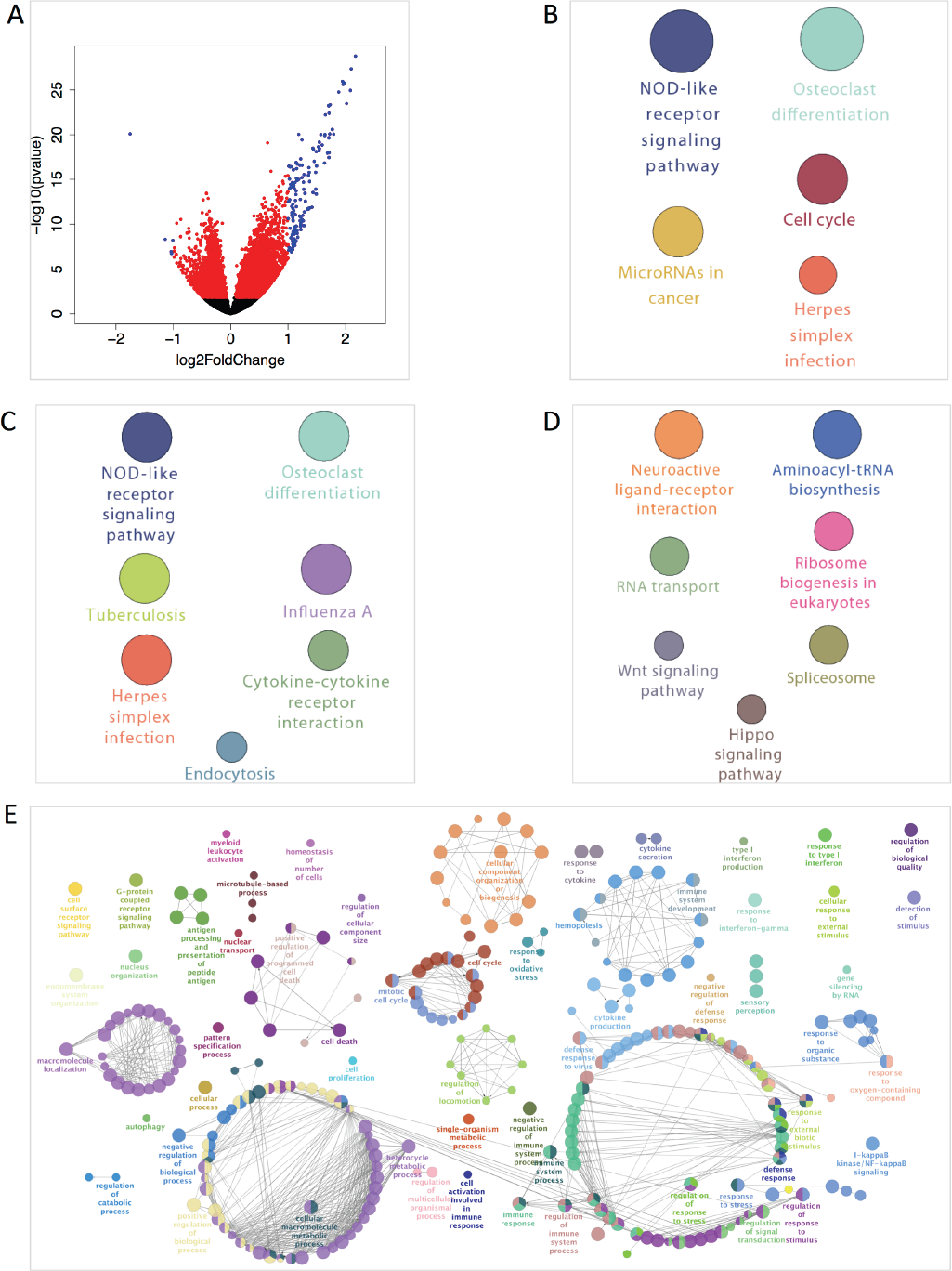
**(A)** Volcano plot of DEGs between SLE patients and healthy individuals. X-axis indicate the log2 fold change (SLE/healthy) while the y-axis indicates the –log10 of the Benjamini-Hochberg adjusted p-value. Black dots indicate genes that do not pass the 5% FDR threshold of significance, red dots indicate genes that do pass the 5% FDR threshold of significance while blue dots indicate genes that pass the significance threshold and have log2fold change >1. The up-regulated genes in SLE are more (3977) compared to the down regulated (2753) and show higher significance. **(B)** Pathway enrichment analysis with all the DEGs. Each circle represents a significantly enriched pathway. The size of the circle represents higher enrichment of the specific pathway. The enrichment p-values were calculated by a two-sided hypergeometric test and corrected for multiple testing with Bonferroni. The threshold for Bonferroni corrected p-values was set to 0.05. **(C)** Pathway enrichment analysis for the up-regulated in SLE genes. **(D)** Pathway enrichment analysis for the down-regulated in SLE genes. **(E)** Mechanistic map of the biological regulation in SLE. For all the DEGs functionally grouped networks of enriched GO term categories were generated. Each node represents a GO term and the size of each node represents the enrichment significance. The connection between nodes is based on Cohen’s kappa statistic score (≥ 0.4) which depends on the gene sharing between nodes. For each network, only the most significant node is labeled.

**Supplementary Figure 3:**
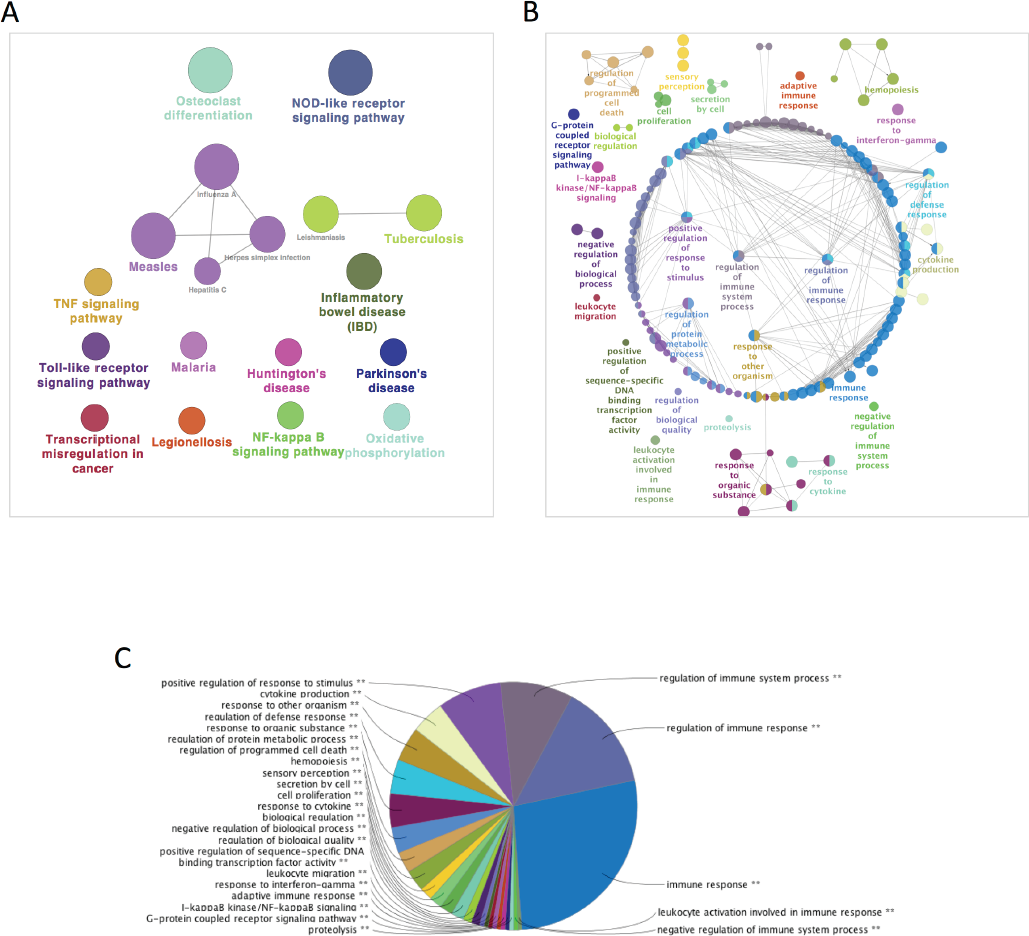
(**A**) Pathway enrichment analysis of the 2726 DEGs (5% FDR) defining the core disease signature. Each circle represents a significantly enriched pathway. The size of the circle represents higher enrichment of the specific pathway. The enrichment p-values were calculated by a two-sided hypergeometric test and corrected for multiple testing with Benjamini-Hochberg. The threshold for the corrected p-values was set to 0.05 (5% FDR). **(B)** Mechanistic map of the biological regulation of the core SLE signature. Functionally grouped networks of enriched GO term categories were generated. Each node represents a GO term and the size of each node represents the enrichment significance. The connection between nodes is based on Cohen’s kappa statistic score (≥ 0.4) that depends on the gene sharing between nodes. For each network, only the most significant node is labeled. **(C)** Pie chart representation of the mechanistic map. More than half of the networks are related to immunity response, system and regulation.

**Supplementary Figure 4:**
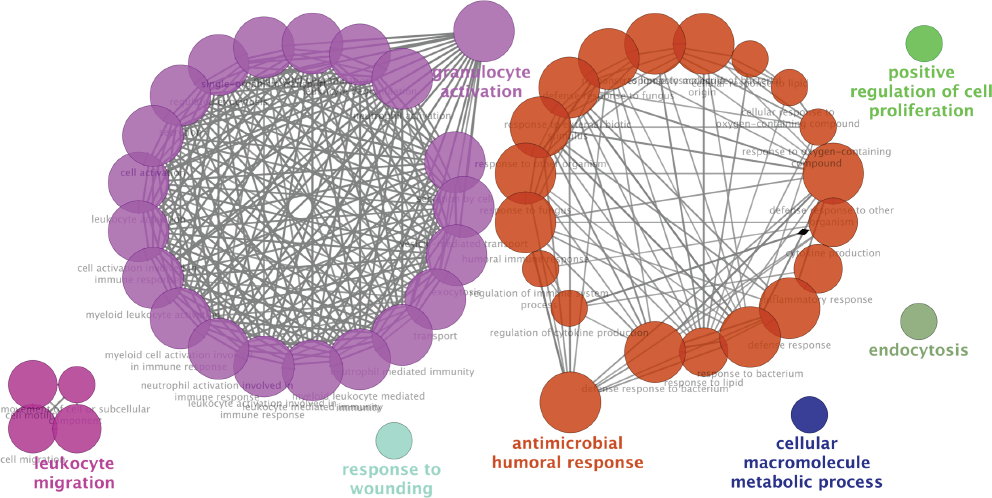
Functionally grouped networks of enriched GO term categories were generated for the 136 DEGs (5% FDR) between group 1 (renal activity) and groups 2,3,4 (activity from other organs). The main enriched terms are granulocyte activation and antimicrobial humoral response supporting the findings that Neutrophils plays crucial role in Lupus Nephritis pathogenesis.

**Supplementary Figure 5:**
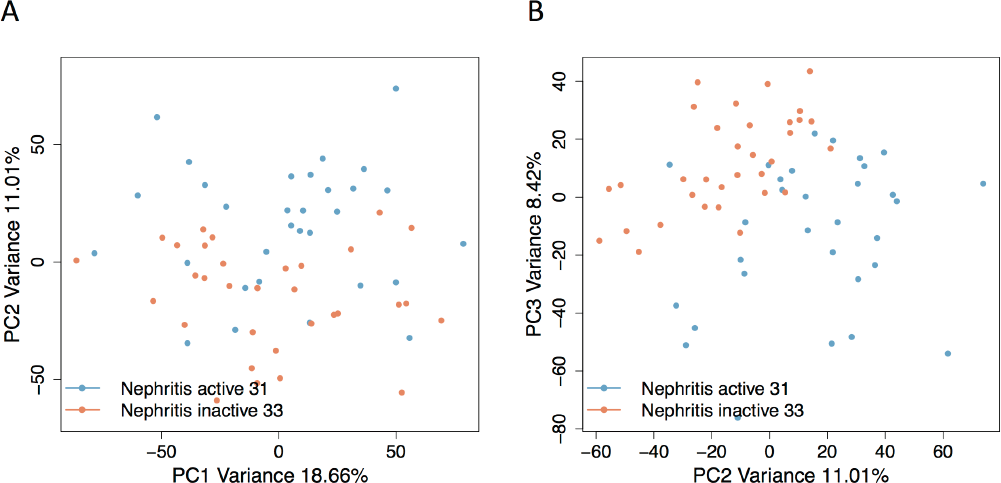
PCA analysis of gene expression between active Nephritis and Nephritis in remission. **(A)** PC1 and PC2 are plotted in x and y-axis respectively. PC2 is differentiating the 2 groups. **(B)**PC2 and PC3 are plotted in x and y-axis respectively. In the differentiation PC3 is participating implying different biological aberrations captured by PC2 and PC3.

**Supplementary Figure 6:**
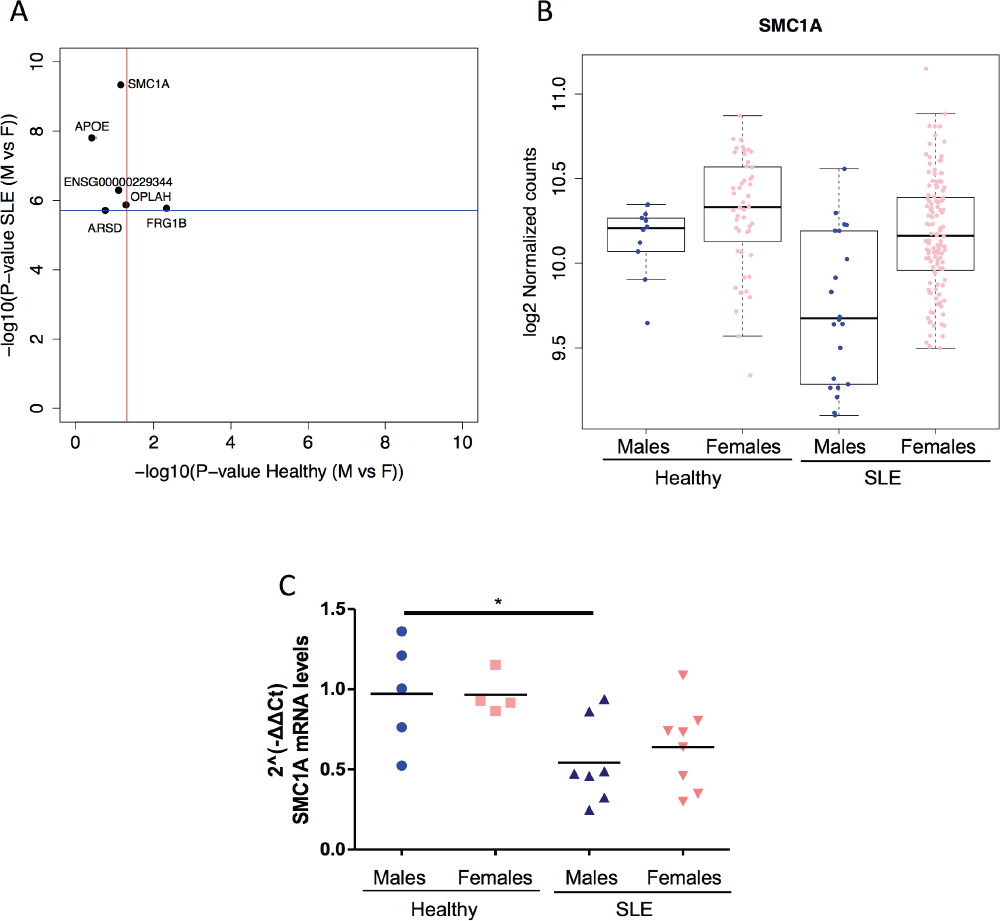
**(A)** Genes with differential expression (Bonferroni significant) between male and female SLE patients that are not differentially expressed (90% FDR) between male and female healthy individuals. The blue line indicates the Bonferroni p-value threshold while the red line is set at p-value 0.05. (**B)** SMC1A expression (normalized RNA-seq) levels in male and female SLE and healthy individuals. **(C)** Scatter plot with bar of SMC1A mRNA expression in purified CD4+ T-cells from the peripheral blood of male and female SLE and healthy individuals. Bars represent median values (two-way ANOVA [F (1, 20) = 11.48, p-value = 0.0029] followed by Tukey’s multiple comparisons test; * p-value <0.05 for the comparison between healthy and SLE males).

**Supplementary Figure 7:**
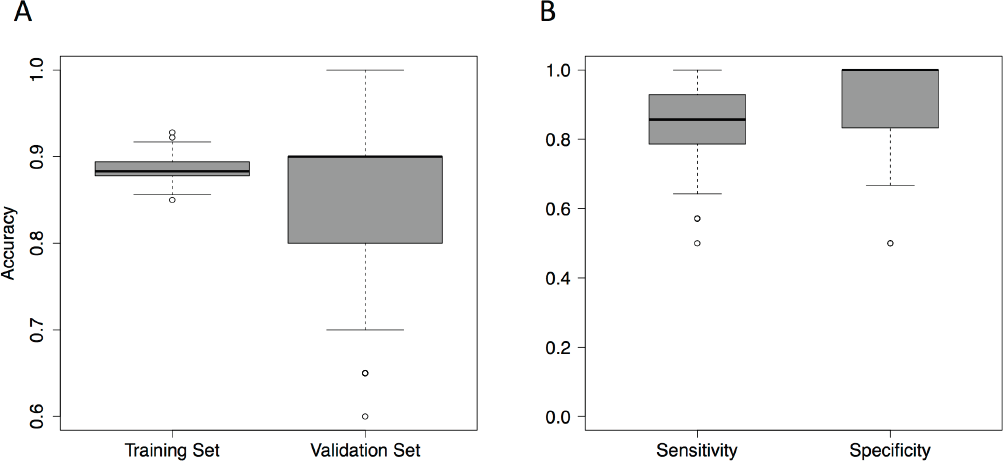
**(A)** Distribution of accuracy after performing 1000 iterations for the training and validation set. Median accuracy of the validation set is 0.9 while for the training set 0.88. **(B)** Distribution of sensitivity and specificity of the 1000 LDA classifiers. Median sensitivity is 0.86 while median specificity is 1.

**Supplementary Figure 8:**
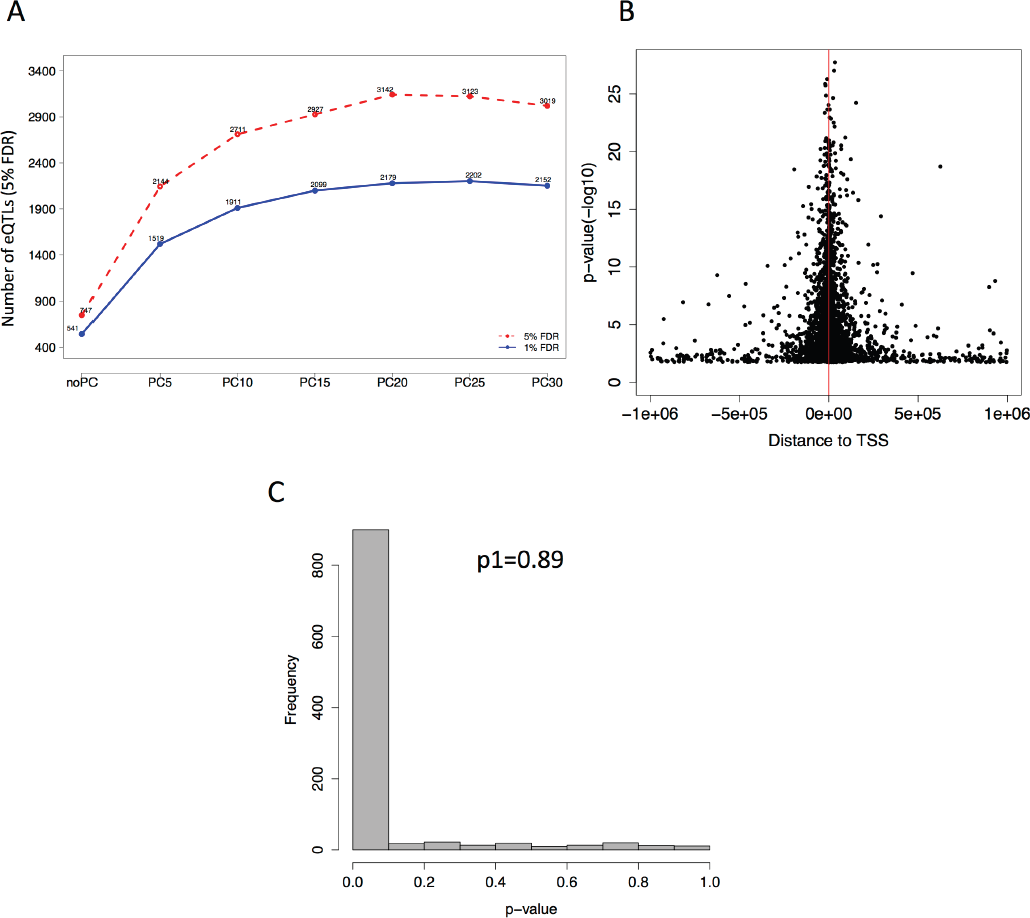
**(A)** Number of detected cis-eQTLs using 142 SLE individuals at 5% FDR (red dotted line) and at 1% FDR (blue continues line) without correction and by using different number of PCs. We detect the higher number of eQTLs (3142 5% FDR) by correcting for the first 20 PCs. **(B)** Distance of cis-eQTLs from the Transcriptional Start Site (TTS) of the genes. The significance of the associations is plotted in the y-axis (-log10 p-value). Highly significant eQTLs are clustered close to TSS. **(C)** Replication of significant eQTLs identified in SLE in another whole blood healthy cohort.

